# MEKK3-MEK5-ERK5 Signaling Promotes Mitochondrial Degradation

**DOI:** 10.1101/2020.05.15.098442

**Authors:** Jane E. Craig, Joseph N. Miller, Raju R. Rayavarapu, Zhenya Hong, Gamze B. Bulut, Wei Zhuang, Sadie Miki Sakurada, Jamshid Temirov, Jonathan A. Low, Taosheng Chen, Shondra M. Pruett-Miller, Lily Jun-shen Huang, Malia B. Potts

## Abstract

Mitochondria are vital organelles that coordinate cellular energy homeostasis and have important roles in cell death. Therefore, the removal of damaged or excessive mitochondria is critical for maintaining proper cellular function. The PINK1-Parkin pathway removes acutely damaged mitochondria through a well-characterized mitophagy pathway, but basal mitochondrial turnover occurs via distinct and less well-understood mechanisms. Here we report that the MEKK3-MEK5-ERK5 kinase cascade is required for mitochondrial degradation in the absence of exogenous damage. We demonstrate that genetic or pharmacological inhibition of the MEKK3-MEK5-ERK5 pathway increases mitochondrial content by reducing lysosome-mediated degradation of mitochondria under basal conditions. We show that the MEKK3-MEK5-ERK5 pathway plays a selective role in basal mitochondrial degradation but is not required for non-selective bulk autophagy, damage-induced mitophagy, or restraint of mitochondrial biogenesis. This illuminates the MEKK3-MEK5-ERK5 pathway as a positive regulator of mitochondrial degradation that acts independently of exogenous mitochondrial stressors.

## Introduction

Maintaining a population of healthy mitochondria is vital for proper development and is necessary to preserve tissue and organ function in response to stress conditions (Pickles, Vigie, & Youle, 2018). Mitochondria are well known for their importance in metabolism and energy production, but they are also a major source of reactive oxygen species (ROS), cytosolic DNA, and pro-apoptotic proteins (Kausar, Wang, & Cui, 2018; McArthur et al., 2018; Murphy, 2009; West et al., 2015). Therefore, it is important to maintain sufficient quantities of healthy mitochondria to meet current metabolic demand, but dangerous to carry an excessive surplus of these organelles. Damaged or dysfunctional mitochondria tend to generate higher levels of ROS and cytosolic DNA compared to healthy mitochondria (Kausar et al., 2018; Murphy, 2009; Sliter et al., 2018). Exposure to high levels of ROS can cause DNA mutations in both nuclear and mitochondrial DNA and can induce protein misfolding throughout the cell (Kausar et al., 2018; Murphy, 2009). Mutations in mitochondrial DNA further decrease the health of the mitochondrial population and promote additional ROS production (Hahn & Zuryn, 2019; Mattiazzi et al., 2004; Nissanka & Moraes, 2018). Thus to maintain homeostasis, the cell has evolved complex systems for the quality control of mitochondria (Ding & Yin, 2012). These systems are designed to balance the elimination of dysfunctional or superfluous mitochondria with mitochondrial biogenesis (Pickles et al., 2018). There are certain situations in which mitochondrial content changes even in the absence of damage, such as during differentiation of highly specialized cell types or induction of a metabolic shift towards a glycolytic phenotype or an oxidative phosphorylation phenotype (Esteban-Martinez & Boya, 2018; Esteban-Martinez et al., 2017; Hu et al., 2016; Palikaras, Lionaki, & Tavernarakis, 2018).

A major mechanism of mitochondrial degradation is known as mitophagy, a form of selective autophagy (Ding & Yin, 2012). Autophagy is an evolutionarily conserved process that uses double membrane vesicles known as autophagosomes to sequester cellular components like damaged or excessive organelles and protein aggregates (Mizushima, 2007). Autophagosomes then fuse with highly acidic lysosomes that contain hydrolases which degrade the components inside (Dikic, 2017; Mizushima, 2007). Mitophagy follows the same steps as autophagy, but the autophagosome selectively forms around mitochondria that have been tagged for degradation by selective autophagy receptors such as NIX, OPTN, or p62/SQSTM1 (Ding & Yin, 2012). These proteins function by binding both the cargo and ATG8-family proteins (commonly referred to as “LC3”) that decorate the forming autophagosome (Yamada, Dawson, Yanagawa, Iijima, & Sesaki, 2019).

Mitophagy plays important roles in development and in disease prevention. Insufficient mitochondrial clearance impairs erythroid maturation and can result in anemia (Sandoval et al., 2008). Removal of paternal mitochondria after zygote formation increases fitness, as mitochondrial DNA is at increased risk of ROS damage during the process of spermatogenesis and fertilization (Sutovsky et al., 2000). Parkinson’s disease, the second most common neurodegenerative disorder, is characterized by dysfunctional mitochondria and compromised mitophagy within the affected neurons (Gao et al., 2017; Kalia & Lang, 2015; Narendra, Tanaka, Suen, & Youle, 2008). Additionally, dysfunctional mitochondria can contribute to the metabolic reprogramming of cancer cells (Gaude & Frezza, 2014). Understanding how novel regulatory pathways contribute to the maintenance of mitochondrial homeostasis in different tissues could provide insights into the development of targeted treatments when these systems fail in disease.

Early insights into the mechanistic basis of mitochondrial degradation arose from studies in yeast, but how mitochondrial degradation is controlled in multicellular organisms such as humans is not yet fully understood (Chen & Klionsky, 2011; Glick, Barth, & Macleod, 2010; Nakatogawa, Suzuki, Kamada, & Ohsumi, 2009). Pink1-Parkin is the most well studied mitophagy regulatory pathway in animals. Though it has been very well characterized, most studies depend on ectopic Parkin overexpression and mitochondrial depolarization with CCCP or other pharmacological agents (Drake, Springer, Poole, Kim, & Macleod, 2017; Lazarou et al., 2015; Narendra et al., 2008). Endogenous Parkin is poorly expressed in several mammalian tissues and cell lines, and it is clear that mitophagy can occur independently of Parkin and PINK1 (Huynh, Dy, Nguyen, Kiehl, & Pulst, 2001; McWilliams et al., 2018).

Here, we report the identification of a previously undescribed role for the MEKK3-MEK5 signaling pathway as a regulator of basal mitophagy in cultured mammalian cells. MEKK3 and MEK5 are upstream activators of the extracellular signal-regulated kinase 5 (ERK5), which is the most recently discovered member of the MAPK family of proteins (Nithianandarajah-Jones, Wilm, Goldring, Muller, & Cross, 2012). ERK5 is ubiquitously expressed in mammalian tissues and is activated by extracellular signals such as growth factors, several cellular stressors, and oxidative phosphorylation (Khan et al., 2018; Nishimoto & Nishida, 2006; Nithianandarajah-Jones et al., 2012). Mouse genetic studies have demonstrated that MEKK2/3-MEK5-ERK5 signaling is required for early embryogenesis, development of the vasculature and muscles, endothelial cell function and cardio protection (Hayashi et al., 2004; W. Liu et al., 2017; Nishimoto & Nishida, 2006; Nithianandarajah-Jones et al., 2012). Alterations in the MEKK2/3-MEK5-ERK5 pathway have been associated with several human diseases and disorders including cancer, childhood obesity, scoliosis, and cerebral cavernous malformation (Cullere, Plovie, Bennett, MacRae, & Mayadas, 2015; Simoes, Rodrigues, & Borralho, 2016; Zhou et al., 2018; Zhu et al., 2014). Our studies indicate that MEKK3-MEK5-ERK5 signaling is critical for basal mitochondrial degradation in mammalian cells and plays a relevant role in erythrocyte maturation.

## Materials and Methods

### Cell Culture

Transformed MEFs, U2OS cells stably expressing mitochondrial targeted mCherry, U2OS cells stably expressing GFP-LC3B, and HeLa YFP-Parkin cells were received from Michael White’s laboratory. The U2OS GFP-LC3B cells were originally a kind gift from Xiaodong Wang, and the HeLa YFP-Parkin cells were originally a kind gift from Richard Youle. Mito-mKeima stably expressing U2OS cells were generated by transfection (addgene #56018), selection with 1mg/ml G418 for 2 weeks, and flow sorted for high expressers. Cells were grown in DMEM supplemented with 10% FBS, L-Glutamine, Penicillin and Streptomycin, and Sodium Pyruvate, and at 37° C and 5% CO_2_. Mito-mCherry, GFP-LC3B, and mito-mKeima stable cells were maintained in 1 mg/ml G418 DMEM media.

### CRISPR/Cas-9 knockout cell lines

SQSTM1/p62 and MAPK7/ERK5 knockout U2OS cells were generated using CRISPR-Cas9 technology. Briefly, 400,000 U2OS cells were transiently transfected with precomplexed ribonuclear proteins (RNPs) consisting of 100 pmol of chemically modified sgRNA (sgSQSTM1 5’ - gagggaaagggcuugcaccg - 3’ or sgMAPK7 5’ – agacggcgaggacggcucug - 3’, Synthego) 35 pmol of Cas9 protein (St. Jude Protein Production Core), and 200ng of pMaxGFP (Lonza) via nucleofection (Lonza, 4D-Nucleofector™ X-unit) using solution P3 and program CM-104 in a small (20ul) cuvette according to the manufacturer’s recommended protocol.

Five days post nucleofection, cells were single-cell sorted by FACS to enrich for GFP+ (transfected) cells, clonally selected, and verified for the desired targeted modification via targeted deep sequencing using gene specific primers with partial Illumina adapter overhangs (hSQSTM1.F – 5’ - acagccccacagtgacgacagaggg - 3’ and hSQSTM1.R 5’- tgggggaggaattagcagagcggca - 3’ or hMAPK7.F – 5’ agctagtctgccacgaaccagccgc 3’ and hMAPK7.R – 5’ cgggcggaggacaccactccatagg 3’, overhangs not shown). NGS analysis of clones was performed using CRIS.py (Connelly & Pruett-Miller, 2019). Genotypes of each clone are available upon request.

### Inhibitor treatments

Unless otherwise stated cells were treated with 10 µM BIX02189 (Tocris 4842) and 10 µM XMD8-92 (Tocris 4132) for 16 hours and 50 nM Bafilomycin A1 (Sigma B1793) for 2 hours.

### Immunoblotting

Western blots were performed by lysing cells in 2x Laemmli loading buffer with 10 mM DTT, separated on polyacrylamide gels, transferred to PVDF membranes and probed with antibodies against p62/SQSTM1 (CS 51145), XPB (SC-293), TOMM40 (MBL 3740), B-actin (Sigma A1978), VDAC (CS 4867S), S6K (CS 9202S), ERK5 (CS 3372S), mtCOX2 (Ab797393), MEKK3 (SC-28769), MEK5 (Invitrogen PA5-15083), PGC1a (CS 2178S), pERK5 (CS 3371S).

### Mitochondrial Content

Relative mitochondrial content was assessed either qualitatively by western blotting for endogenous mitochondrial proteins and non-mitochondrial loading controls (described above) or via quantitative analysis of fluorescently-labeled mitochondria via flow cytometry. Mitochondria were fluorescently labeled either by stable expression of mitochondrial targeted mCherry (mito-mCherry) or staining with Mito-Tracker Green-FM (Thermo Fisher, M7514). U2OS cells stably expressing mito-mCherry were plated in 6-well plates for flow cytometry analysis or in Nunc Lab-Tek II chambered coverglass (Thermo, 155382) for microscopy imaging. Alternatively, cells were stained with 100 nM Mito-Tracker Green FM for 30 minutes then washed with PBS three times. Cells were then trypsinized and assayed as described in the Flow Cytometry section below. Mitochondrial content per cell was measured from 10,000 cells per condition by quantifying the fluorescence of either mito-mCherry or Mito-Tracker Green FM, and the median value for each condition was normalized to the median value from the appropriate control condition.

### Mitochondrial uncoupling

HeLa YFP-Parkin cells were treated with 10 nM CCCP overnight in addition to indicated treatments. Cells were then fixed and stained with DAPI and TOMM20 antibody.

### Confocal Imaging

Regular confocal images were taken with Leica SP8 TCS equipped with White Light Laser, 405 nm diode laser using 63x/1.4NA/Oil/HC PL APO CS2/0.14mm objective confocal microscope. Mito-mCherry images were created from Z-stacks taken at software calculated Nyquest pixel resolution. Higher resolution confocal images were taken using 100x/1.4NA objective with oversampled XY pixel size of 16 nm and Z-step size of 100 nm which were deconvolved using Huygens Professional 16.10 (Scientific Volume Imaging, Netherlands).

GFP-LC3B puncta and SQSTM1/p62 puncta were manually counted on a per cell basis.

MAPK7/ERK5 wild-type and MAPK7/ERK5 knockout U2OS cells stably expressing mitochondrial targeted mKeima were plated in Nunc Lab-Tek II chambered coverglass and imaged live on the Leica SP8 TCS confocal microscope. Data were analyzed by comparing 561 (acidic) signal to 458 (neutral) signal.

### Lysosomal Acidification

Lysosomal content was determined by staining with LysoTracker Red DND-99 (L7528) at 100nM for 30 minutes. Lysosomal acidification was determined by staining with LysoSensor Green DND-189 (L7528) at 1mM for 10 minutes. Cells were then imaged using a BioTek Cytation 5 Cell Imaging Multi-Mode Reader. Cells were imaged using 20x objective. Signal was analyzed via relative fluorescence using CellProfiler (McQuin et al., 2018).

### Flow cytometry

Cells were treated as indicated, trypsinized, spun down at 400 rpm, and resuspended in buffer made from 1 x PBS, 2% serum, 1 mM EDTA, and 0.1% Sodum Azide. Cells were stained with DAPI. Fluorescence was then quantified on a BD Biosciences Fortessa by gating on single live cells and subsequently analyzed using FloJo v10.6.2.

### RNAseq

RNA was extracted from cultured cells using RNeasy Kit (QIAGEN #74104) following manufacturers’ protocols. RNA concentration was measured using a NanoDrop (Thermo Fisher scientific, Waltham, MA) and the quality of RNA was determined with a bioanalyzer (Agilent Technologies, Santa Clara, CA). Libraries were prepared using the TruSeq Stranded Total RNA Library Prep Kit (Illumina, San Diego, CA) and subjected to 100 cycle paired-end sequencing on the Illumina HiSeq platform.

### Promega Autophagy Assay

HEK293 autophagy reporter cells (HiBiT-HaloTag-LC3, Promega #GA1040) were grown in in DMEM (ThermoFisher #31053-028) supplemented with 10% FBS (Hyclone #SH30071), L-Glutamine (ThermoFisher #25030-081), Penicillin and Streptomycin (ThermoFisher #15140-122), Sodium Pyruvate (ThermoFisher #11360-070), and 500 µg/mL G418 (ThermoFisher #10131) at 37° C and 5% CO_2_. 2500 cells in 25 µl of G418-free media were plated into each well of Corning 8804BC 384-well plates and grown for 18 hours. Cells were then treated with 10-point curves with 3-fold dilutions and a top concentration of 60 µM of Bafilomycin A1 (positive control for autophagy inhibition), Rapamycin (TOCRIS #1292; positive control for autophagy activation), BIX02188, BIX02189, and XMD8-92, or DMSO using a V&P Scientific S100 pin-tool before they were returned to 37°C with 5% CO_2_ for 24 hours. The assay plates were then equilibrated to room temperature for 15 minutes before the addition of 25 µl of Nano-Glo HiBiT Lytic Detection System (Promega #N3030.) The plates were then shaken at 300 rpm for 2 minutes on an orbital shaker, incubated for 30 minutes at room temperature, and the luminescence of each well measured using a Perkin Elmer Envision plate reader.

### Erythroblast maturation

Ter119-negative erythroid progenitors were isolated from embryonic day 13.5 or 14.5 Balb/c murine fetal livers as described (Sulahian, Cleaver, & Huang, 2009). The purity of the cells was confirmed by staining cells with PE-CD71 and allophycocyanin (APC)-Ter119 antibodies (all from BD Biosciences) by flow cytometry. Approximately 10% of total fetal livers were purified as Ter119 negative. These progenitors were set up on retronectin coated plates for differentiation and were followed over three days for differentiation status and mitochondrial content. The data were acquired on a BD FACSCalibur and analyzed by FlowJo 8.8.4. To measure mitochondrial content, cells were stained with 25nM MitoTracker Deep Red FM (Molecular Probes, Invitrogen) for 30 min at 37°C.

For inhibitor treatments starting on Day 0, Ter119-negative progenitors were set up for differentiation directly after purification in erythropoietin (Epo) media in the presence of inhibitor on retronectin coated plates. For inhibitor treatments starting on Day 2 and Day 3 Ter119-negative progenitors, 20 hours after culturing in Epo medium or 1 day after removal from EPO medium, were set up for differentiation separately, and inhibitors were added when indicated. Erythroid differentiation was monitored on days 1, 2 and 3 with PE-CD71 (BD Biosciences) and APC-Ter119 (eBioscience) by flow cytometry. 7-AAD (BD Biosciences) staining was used to exclude dead cells. Cells were treated with DMSO, 10 µM BIX02188, or 10 µm XMD8-92.

## Results

### SQSTM1/p62 Constitutively Delivers Mitochondria to Lysosomes for Degradation in the Absence of Exogenous Damage

The primary question we set out to address is what controls basal mitochondrial degradation in the absence of exogenous damage. This process is known to be mechanistically distinct from damage-induced mitophagy, which requires the kinase PINK1, the E3 ligase Parkin, and the selective autophagy receptors NDP52 and Optineurin (Lazarou et al., 2015; McWilliams et al., 2018). We first investigated whether an alternative selective autophagy receptor, SQSTM1/p62, supports mitochondrial degradation in the absence of exogenous damage. We selected p62 as an initial candidate because it cannot support damage-induced, Parkin-dependent mitophagy but has been reported to be important for mitochondrial degradation in other contexts (Lazarou et al., 2015; Matsumoto, Shimogori, Hattori, & Nukina, 2015; Nguyen et al., 2019; Yamada et al., 2019; Yamada et al., 2018). We treated U2OS osteosarcoma cells with Bafilomycin A1 for two hours to inhibit degradation of all cargo delivered to the lysosome and used immunofluorescent microscopy to determine whether such cargo included mitochondria and p62. Our experiments revealed that a minority of the mitochondrial population (marked by TOMM20) were delivered to LAMP1-positive lysosomes in the absence of any mitophagy-inducing treatment, and all LAMP1-encapsulated mitochondria were co-labeled with p62 (Fig. 1A and 1B). In contrast, the vast majority of LAMP1-negative mitochondria were negative for p62 (Fig. 1A and 1B). In the absence of Bafilomycin treatment we observed rare instances of p62-positive mitochondria co-localized with LAMP1 (Fig. 1A).

**Figure 1:**
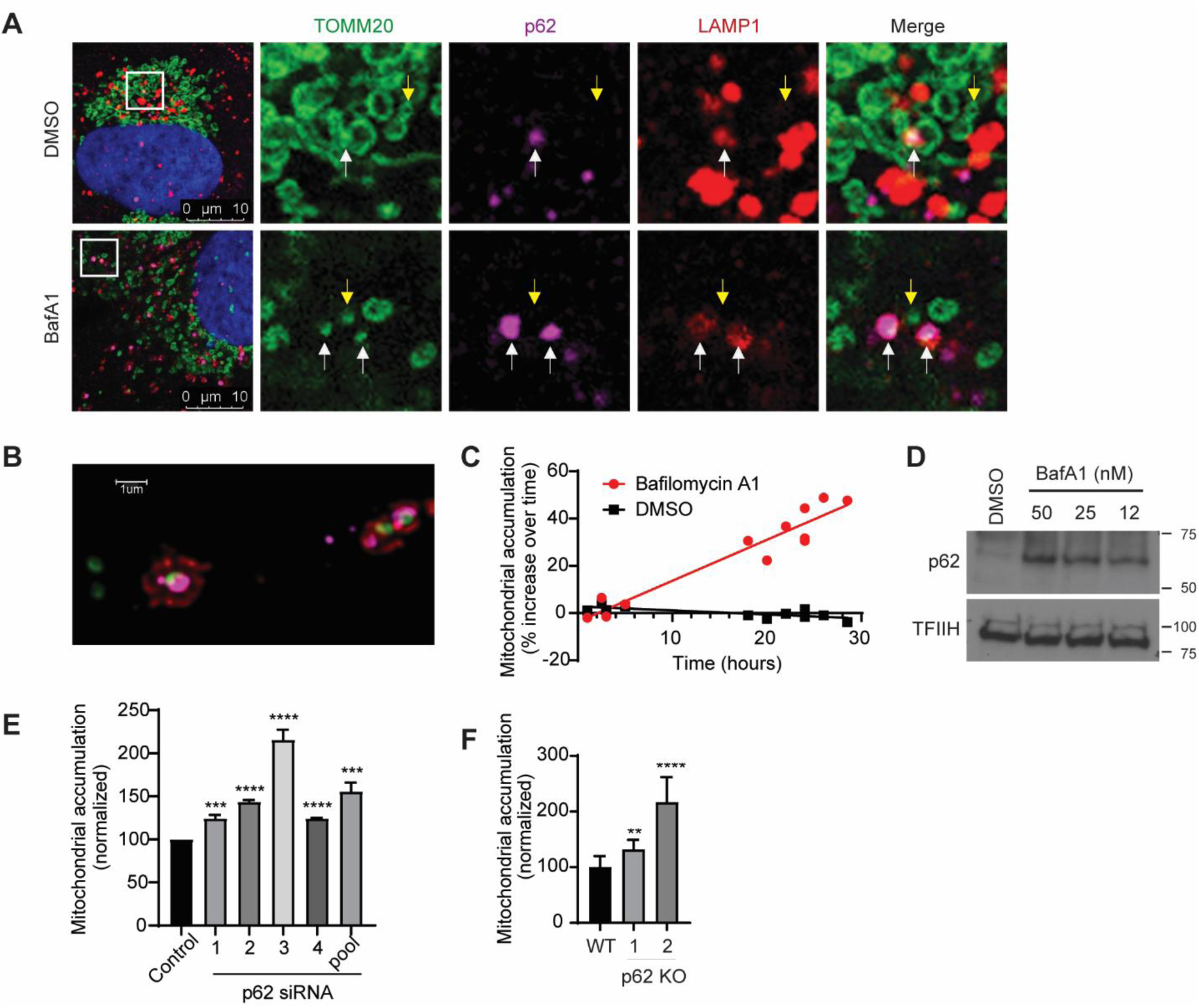
SQSTM1/p62 Constitutively Delivers Mitochondria to Lysosomes for Degradation in the Absence of Exogenous Damage. (A) U2OS cells were treated with vehicle (DMSO) or with 50 nM Bafilomycin A1 for 2 hours to block degradation of lysosomal contents and analyzed by immunocytochemistry. Representative images are shown. TOMM20 (Green), p62 (purple), LAMP1 (red), Dapi (blue). Images show instances of mitochondria (TOMM20) tagged by p62 and surrounded by lysosomes (LAMP1) designated by white arrows and mitochondria that are not being degraded designated by yellow arrows. (B) Representative high-resolution confocal image of cells treated with bafilomycin A1 as in (A). Note LAMP1-positive lysosomal membrane surrounding p62-tagged mitochondria. Tomm20 (green), p62 (purple), LAMP1 (red). (C) U2OS mito-mCherry cells were treated with 50 nM Bafilomycin A1 or vehicle (DMSO) for the indicated time. Mitochondrial accumulation was measured by flow cytometry and displayed as percent increase over time. (D) U2OS cells were treated with the indicated concentrations of Bafilomycin A1 overnight. (E) U2OS mito-mCherry cells were transfected with the indicated siRNA oligos or siControl. (The p62 Si pool contains oligos 1-4). 72 hours later mitochondrial accumulation was measured by flow cytometry. Mean +/- S.E. of n = 3 independent experiments is shown; *** p < 0.001; **** p < 0.0001. (F) p62 knockout U2OS cells were generated via CRISPR/Cas9-mediated genome editing. Cells were stained with 100 nM Mitotracker Green FM for 30 minutes and mitochondrial accumulation was measured by flow cytometry. Mean +/- S.E. of n = 7 independent experiments is shown; ** p < 0.01; **** p < 0.0001.

To determine whether lysosomal activity detectably limited mitochondrial accumulation under basal conditions, we stably expressed a construct encoding the CoxVIII leader sequence fused to mCherry in U2OS cells (U2OS mito-mCherry cells) and measured mCherry fluorescent intensity per cell by flow cytometry as a marker of mitochondrial abundance. Short durations of lysosomal inhibition (< 5 hours) did not detectably increase bulk mitochondrial content per cell, while longer durations of lysosomal inhibition caused a steady increase of mitochondrial content over time (Fig. 1C). Together these results indicate that steady-state mitochondrial abundance is limited by ongoing delivery of a small subset of p62-labeled mitochondria to lysosomes for degradation. Treatment with Bafilomycin A1 increased total protein levels of p62 in a dose dependent manner, confirming that p62 is continually degraded by the lysosome under basal conditions (Fig. 1D). Together, these results suggest that p62 labels mitochondria destined for delivery to lysosomes in the absence of exogenous damage or ectopic Parkin overexpression.

To test whether p62 is required for basal mitochondrial degradation, we depleted p62 by RNAi or CRISPR/Cas9-mediated gene editing and asked whether total mitochondrial content per cell increased as a result. Multiple independent siRNA oligos targeting p62 caused an increase in mitochondrial accumulation in U2OS mito-mCherry cells (Fig. 1E and Sup. Fig. 1A). Similarly, p62 knockout U2OS cells exhibited increased mitochondrial accumulation relative to parental U2OS cells as measured by Mitotracker Green FM staining (Fig. 1F and Sup. Fig. 1B). Taken together, our data indicate p62 is important for basal delivery of mitochondria to lysosomes for degradation in the absence of exogenous damage.

### Nomination of Putative Mitophagy Regulatory Pathways

To identify novel mitophagy regulatory pathways, we utilized Functional Signature Ontology (FuSiOn), a method for unbiased identification of functionally related proteins, to nominate protein kinases from the human genome that function similarly to p62 under basal conditions (Potts et al., 2013). FuSiOn identified ten kinases (BMP2K, DCLK3, LIMK2, MAP2K5/MEK5, MAP3K3/MEKK3, ROS1, SIK2, TAOK2, ULK1, and ULK2) whose depletion mimicked the phenotypic effect of p62 depletion in HCT116 colon cancer cells (Potts et al., 2013). To determine whether any of these ten candidates promotes basal mitophagy, we depleted each kinase individually in U2OS mito-mCherry cells and measured the resulting change in average mitochondrial content per cell. Depletion of seven of the ten kinases caused an increase in mitochondrial content similar to that caused by depletion of p62 (Fig. 2A). Next we depleted each kinase in U2OS GFP-LC3B cells and measured accumulation of GFP-LC3B as a marker of nonselective autophagy inhibition. Two candidates (MAP3K3 and MAP2K5) phenocopied p62’s selectivity for mitochondria, demonstrated by the relatively consistent LC3B levels in addition to the increase in mitochondrial content (Fig. 2A). MAP3K3 encodes MEKK3, which activates the MEK5-ERK5 kinase cascade. MAP2K5 encodes the MEKK3 substrate MEK5 (Fig. 2B). We confirmed on-target efficacy of the siRNA pools directed against MAP3K3/MEKK3 and MAP2K5/MEK5 (Sup. Fig. 1C). Maximum intensity projection images of U2OS mito-mCherry cells confirmed that the mito-mCherry signal remained localized to mitochondrial network after depletion of MEKK3 or MEK5 and demonstrated the accumulation of mitochondrial network in individual cells, supporting the flow cytometric data (Sup. Fig. 1D).

**Figure 2:**
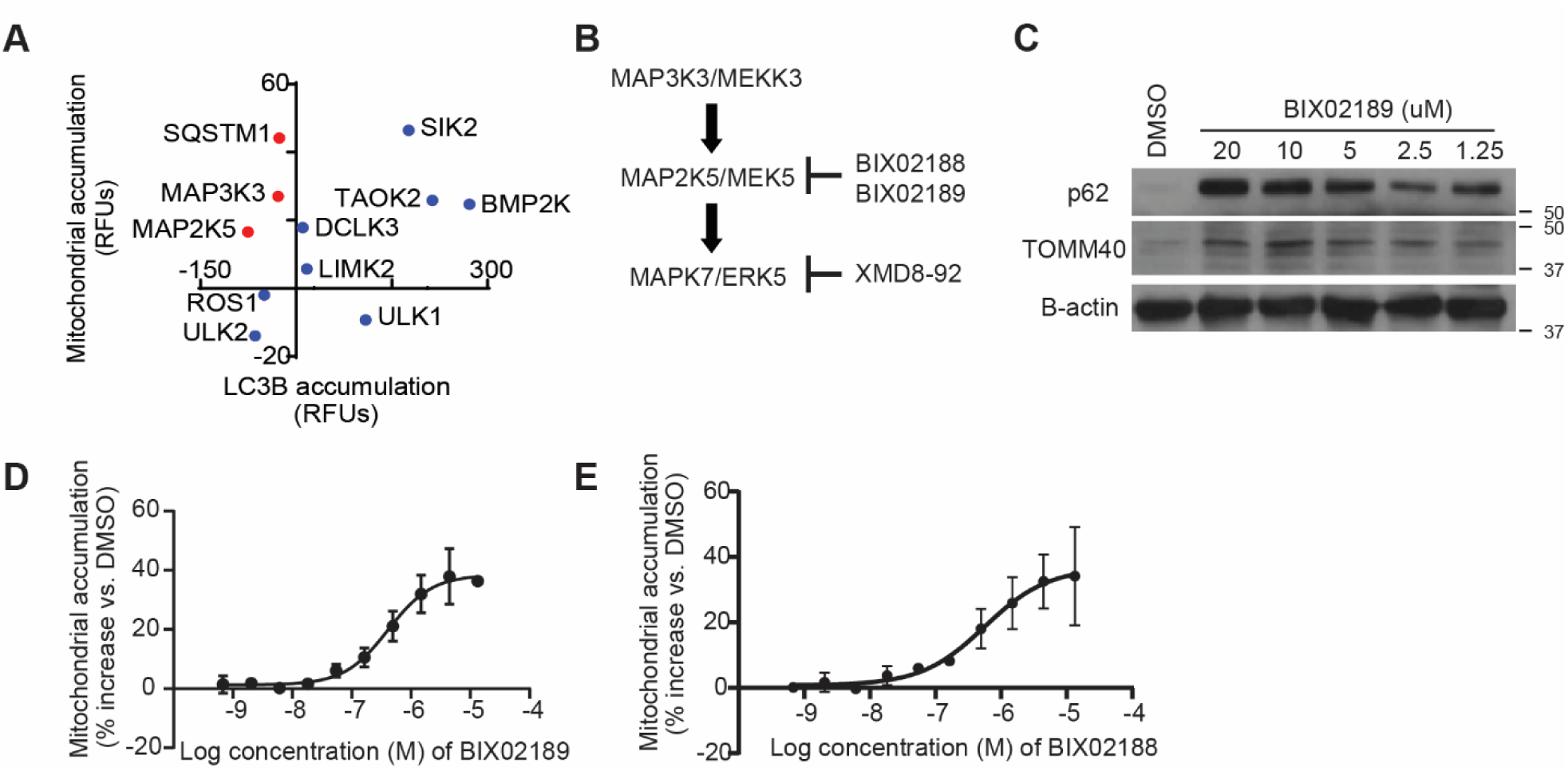
Nomination of Putative Mitophagy Regulatory Pathways. (A) Candidate mitophagy-regulating genes identified by FuSiOn were screened by siRNA-mediated depletion in U2OS mito-mCherry cells and U2OS GFP-LC3B cells. Depletion of 8 of 11 candidates caused mitochondrial accumulation, of which only 3 exhibited selectivity for mitochondria over the bulk autophagy marker GFP-LC3B (red dots). (B) A model of the MAP3K3 kinase cascade including inhibitors. (C) Mouse embryonic fibroblasts were treated with vehicle (DMSO) or the indicated concentration of the MEK5 inhibitor BIX02189 overnight. Lysates were collected and analyzed by western blot using the indicated antibodies. (D) U2OS mito-mCherry cells were treated with vehicle (DMSO) or the indicated concentration of BIX02189 overnight. Mitochondrial accumulation was measured by flow cytometry. Mean +/- S.D. of n = 2 independent experiments is shown. (E) U2OS mito-mCherry cells were treated with vehicle (DMSO) or the indicated concentration of the MEK5 inhibitor BIX02188 overnight. Mitochondrial accumulation was measured by flow cytometry. Mean +/- S.D. of n = 3 independent experiments is shown.

Next, we tested whether pharmacological inhibition of MEK5 kinase activity could alter mitochondrial abundance using two small molecule inhibitors of MEK5 (BIX02188 and BIX02189) (Tatake et al., 2008). Both BIX02188 and BIX02189 inhibited MEK5 activity and increased mitochondrial content in a dose-dependent manner in varied mammalian cells (Fig. 2C-E and Sup. Fig. 1E). Given the structural similarity between the two inhibitors, we measured the off-target activities of BIX02188 and BIX02189 by Kinome Profiling and determined that the off-target activities do not overlap (Sup. Table 1). This increases the likelihood that the observed increase in mitochondrial content is due to inhibition of the intended target, MEK5. Together, these results indicate that MEKK3-MEK5 signaling restrains mitochondrial accumulation.

### The MEKK3-MEK5-ERK5 Kinase Cascade Prevents Accumulation of Excess Mitochondria

We proceeded to assay whether the canonical downstream component of this kinase cascade, ERK5 (encoded by the MAPK7 gene), also plays a role in restraining mitochondrial accumulation. RNAi-mediated depletion of ERK5 increased the average mitochondrial content per cell, indicating that ERK5 (like p62, MEKK3, and MEK5) prevents excess accumulation of mitochondria under basal conditions (Fig. 3A and Sup. Fig. 1D). A small molecule inhibitor of ERK5, XMD8-92 (Yang et al., 2010), also increased mitochondrial content in a dose-dependent manner (Fig. 3B and 3C).

**Figure 3:**
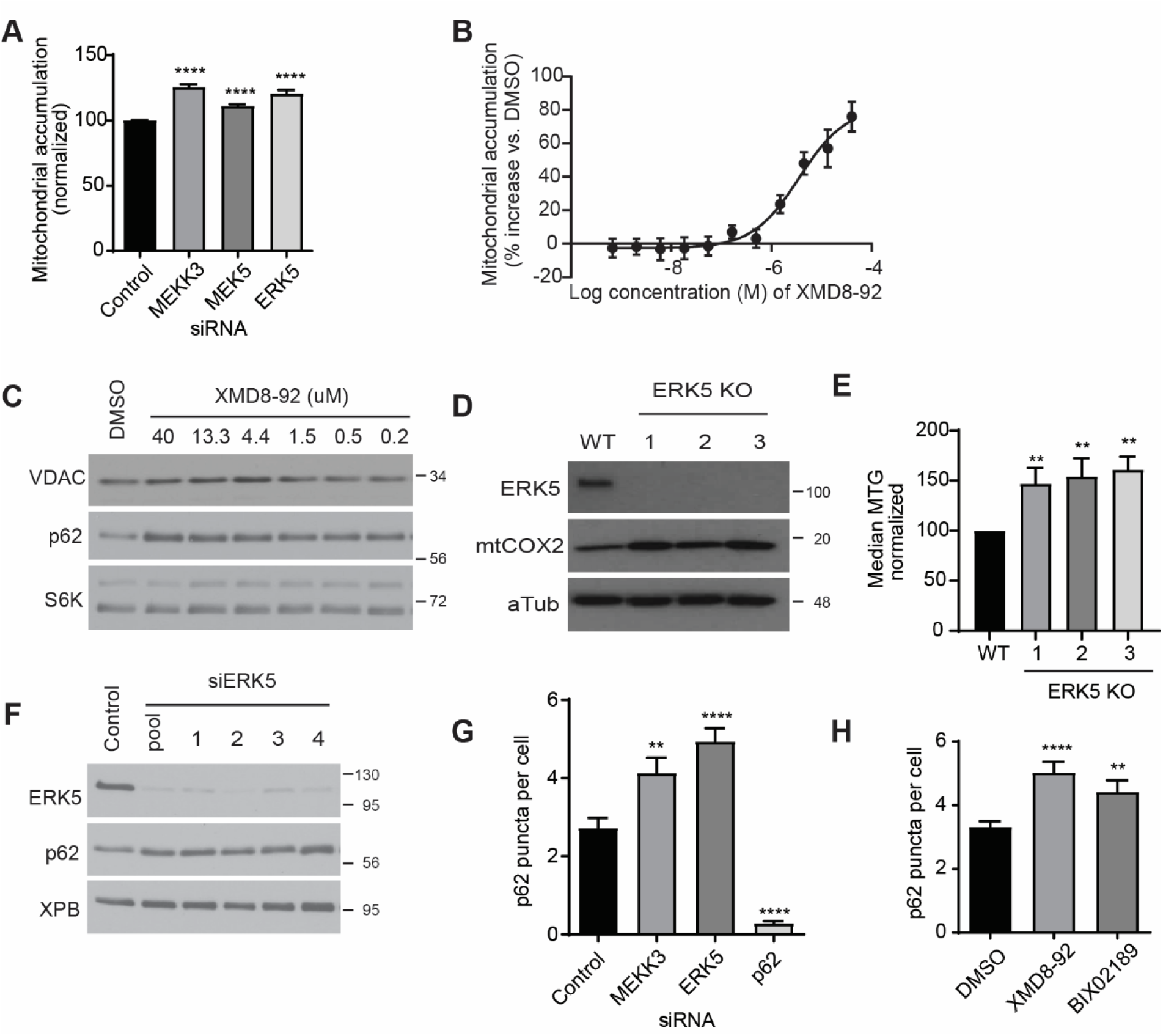
The MEKK3-MEK5-ERK5 Kinase Cascade Prevents Accumulation of Excess Mitochondria. (A) U2OS mito-mCherry cells were transfected with the indicated siRNA oligos. 72 hours later mitochondrial accumulation was analyzed by flow cytometry. Mean +/- S.D. of n = 3 independent experiments is shown. (B,C) U2OS mito-mCherry cells were treated with vehicle (DMSO) or the indicated concentration of XMD8-92 overnight. Mitochondrial accumulation was measured by flow cytometry (B) or western blot (C). Mean +/- S.D. of n = 2 independent experiments is shown (B). (D, E) Three independent MAPK7/ERK5 knockout U2OS clones were generated by CRISPR/Cas9. Mitochondrial levels were assessed by western blot using the indicated antibodies (D) or by Mitotracker Green FM staining and flow cytometry (E). (F) U2OS cells were transfected with the indicated siRNAs. 72 hours later lysates were collected and analyzed by western blot using the indicated antibodies. (G, H) U2OS cells were either transfected with the indicated siRNAs for 72 hours (G) or treated with the indicated drug at 10 µM overnight (H). Cells were fixed and stained with p62 antibody, and then imaged. See Sup. Figure 1 F,G for representative images. Quantitation is shown here.

Thus, we generated MAPK7/ERK5 knockout U2OS cells using CRISPR/Cas9 technology and analyzed mitochondrial content via western blot and MitoTracker Green FM staining. ERK5 knockout cells exhibited increased mitochondrial accumulation relative to the parental controls (Fig. 3D and 3E). These data indicate that the canonical MEKK3-MEK5-ERK5 kinase cascade restrains mitochondrial accumulation under basal conditions.

We asked whether the MEKK3-MEK5-ERK5 kinase cascade promotes mitochondrial degradation through regulation of p62 protein levels. MEKK3, MEK5, and p62 all contain PB1 domains, which mediate protein-protein dimerization (Nakamura, Kimple, Siderovski, & Johnson, 2010). We hypothesized that MEKK3-MEK5-ERK5 pathway inhibition might decrease p62 protein stability, or alternatively, might reduce p62 expression given that ERK5 is known to translocate to the nucleus and regulate gene transcription when activated (Drew, Burow, & Beckman, 2012; Nithianandarajah-Jones et al., 2012). In either of these cases, a decrease in p62 levels upon MEKK3-MEK5-ERK5 pathway inhibition could explain why mitochondria accumulate under those conditions.

However, genetic or pharmacological inactivation of the MEKK3-MEK5-ERK5 kinase cascade induced p62 protein accumulation rather than decrease in abundance, indicating that MEKK3-MEK5-ERK5 signaling is not required for p62 expression or stability (Fig. 2C, 3C, 3F). Using immunofluorescent microscopy, we found that genetic or pharmacological inactivation of the MEKK3-MEK5-ERK5 kinase cascade caused an increase in the average number of cytoplasmic p62 puncta per cell (Fig. 3G and 3H and Sup. Fig. 1F and 1G). The punctate accumulation of the selective autophagy adaptor protein p62 upon inhibition of the MEKK3-MEK5-ERK5 pathway is consistent with the interpretation that this pathway promotes one or more forms of selective autophagy under basal conditions.

### The MEKK3-MEK5-ERK5 Pathway is Required for Lysosomal Degradation of Mitochondria

To determine the underlying cause of increased mitochondrial content observed upon inhibition of MEKK3-MEK5-ERK5 signaling, we considered and tested three distinct possibilities: 1) induction of mitochondrial biogenesis; 2) nonselective inhibition of the autophagy-lysosome pathway; and 3) selective inhibition of mitochondrial degradation. PGC1α is a critical regulator of mitochondrial biogenesis (Jornayvaz & Shulman, 2010), and increased protein levels and activation of PGC1α leads to an increase in transcription factors, TFAM, TFB1M, and TFB2M (Villarroya, Giralt, & Villarroya, 2009). Therefore, if inhibition of MEKK3-MEK5-ERK5 signaling increases mitochondrial abundance by inducing mitochondrial biogenesis, we should detect increased expression of PGC1α, TFAM, TFB1M, and TFB2M upon inhibition of the pathway. RNA-seq confirmed that there was no increase in the mRNA levels of these major mitochondrial biogenesis related transcription factors upon knockout of ERK5 (Fig. 4A, Sup. Table 2). We also verified that the protein levels of the master mitochondrial biogenesis factor PGC1α did not increase upon inhibition of the MEKK3-MEK5-ERK5 pathway. In fact, PGC1α levels decreased upon pharmacological inhibition of MEK5 or ERK5 (Sup. Fig. 2A), consistent with existing literature reporting a positive role for ERK5 signaling in promoting mitochondrial biogenesis (W. Liu et al., 2017). Therefore MEKK3-MEK5-ERK5 signaling does not restrain mitochondrial accumulation by limiting mitochondrial biogenesis.

**Figure 4:**
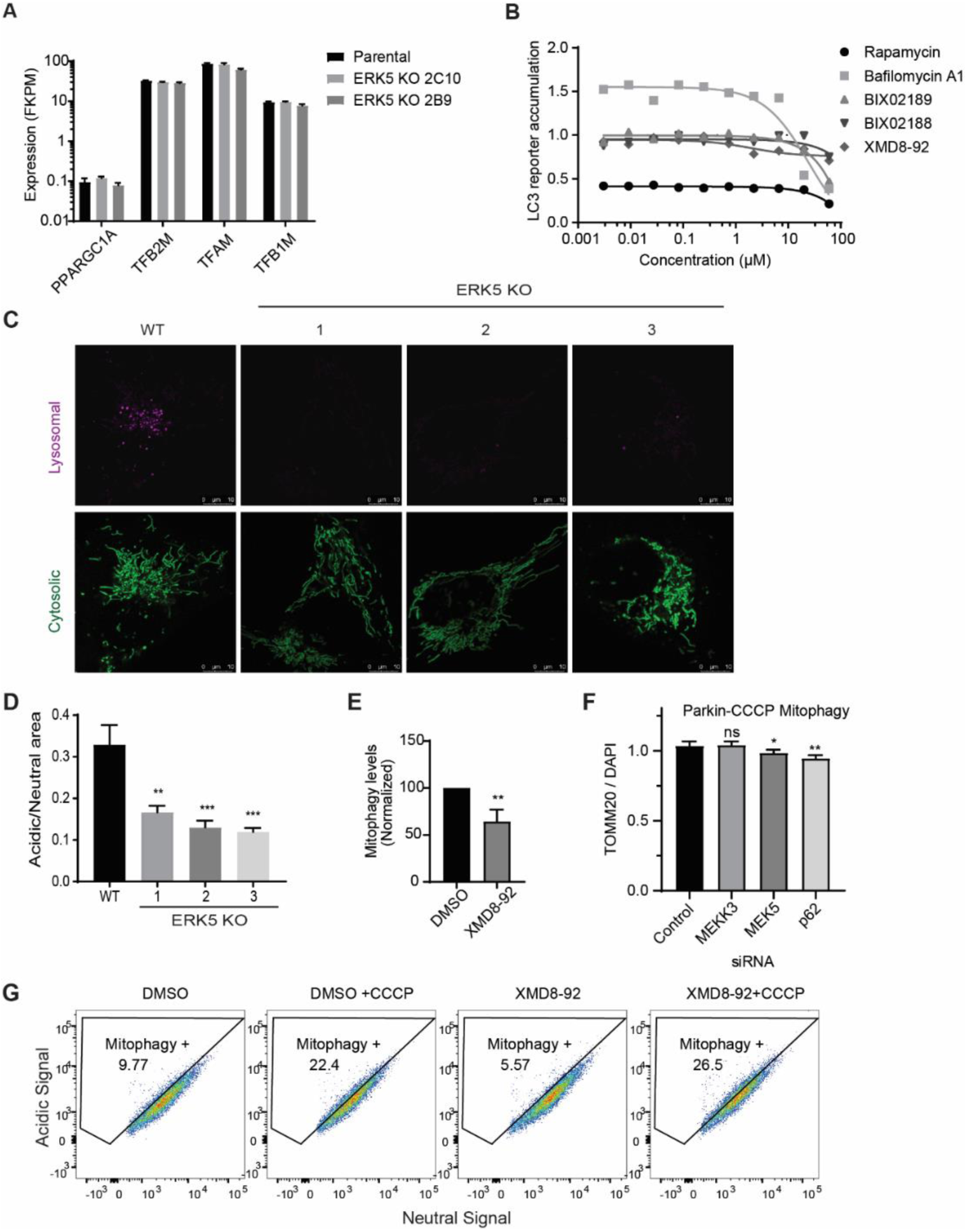
The MEKK3-MEK5-ERK5 pathway is required for lysosomal degradation of mitochondria. (A) RNAseq was performed on parental U2OS cells and two MAPK7/ERK5 knockout U2OS clones. Expression of selected mRNAs encoding mitochondrial biogenesis genes is shown. Note that expression of these genes is not elevated in ERK5 knockout cells relative to parental controls. (B) HEK293 cells stably expressing the HiBiT-LC3 bulk autophagy reporter were treated with vehicle (DMSO) or the indicated compounds at the indicated concentrations for 24 hours in duplicate. Accumulation of the reporter was measured using the Promega Autophagy Assay system and normalized to vehicle-treated cells. (C, D) Parental (WT) and MAPK7/ERK5 knockout U2OS cells were transiently transfected with mitochondrial targeted mKeima. Florescence activated cell sorting was used to select transfection-positive cells which were then analyzed by fluorescent microscopy. Representative images (C) and quantification (D) of active mitophagy events are shown. (E) WT U2OS cells transfected with mitochondrial targeted mKeima were treated for 16 hours with DMSO or 10 µM XMD8-92. Cells were assessed by flow cytometry and mitophagy levels were quantified. Mean +/- S.D. of n = 3 independent experiments is shown. (F) HeLa YFP-Parkin stable cells were transfected with the indicated siRNA for 72 hours and treated with 10 µM CCCP overnight. Cells were fixed, stained with anti-Tom20 and DAPI, and imaged. Mitochondrial content was quantified and normalized to DAPI. (G) WT U2OS cells were stably transfected with mitochondrial targeted mKeima. Cells were treated overnight with vehicle control, 10 µM CCCP, 10 µM XMD8-92 or both. Cells were then collected and analyzed via flow cytometry.

Additionally, MEK5/ERK5 signaling is not required for nonselective autophagy or for general lysosomal function. We performed a Promega Autophagy Assay, in which we treated HEK293 cells expressing the HiBiT-LC3 autophagy reporter with a range of concentrations of DMSO, Bafilomycin A1, Rapamycin, BIX02188, BIX02189, and XMD8-92 for 24 hours. As expected, Bafilomycin A1 effectively inhibited nonselective autophagy and Rapamycin effectively induced autophagy across wide dose ranges, as evidenced by increased (Bafilomycin) or decreased (Rapamycin) accumulation of the HiBiT-LC3 reporter (Fig. 4B). In contrast, BIX02188, BIX02189, and XMD8-92 did not inhibit nonselective autophagy even at concentrations as high as 60 µM (Fig. 4B). Furthermore, we determined that inhibition of the MEK5-ERK5 pathway does not cause lysosomal deacidification or alter lysosomal content (Sup. Fig. 2B and 2C). We also monitored autophagosome formation and autophagic flux using U2OS GFP-LC3B cells and observed no disruptions in either process upon treatment with XMD8-92 (Sup. Fig. 2D and 2E).

The combined findings indicate the MEKK3-MEK5-ERK5 kinase cascade restrains mitochondrial accumulation independent of mitochondrial biogenesis, driving bulk autophagy, or supporting general lysosome function. Next we questioned whether the MEKK3-MEK5-ERK5 kinase cascade is required for selective degradation of mitochondria. Therefore, we developed WT and ERK5 KO U2OS cell lines expressing mitochondrial-targeted monomeric Keima, a pH dependent fluorescent protein (Kogure, Kawano, Abe, & Miyawaki, 2008; Tantama, Hung, & Yellen, 2011) to measure the delivery of mitochondria to lysosomes. When targeted to the mitochondria, the pH dependent fluorescence allows us to distinguish between mitochondria in the cytoplasm (458nm) and mitochondria that are being degraded in the highly acidic lysosome (561nm) (Biel & Rao, 2018). We found that all three independent ERK5 knockout clones had significantly less mitochondria in lysosomes compared to parental cells (Fig. 4C and 4D). Furthermore, in parental U2OS mito-mKeima cells pharmacological inhibition of ERK5 using XMD8-92 decreased the amount of basal mitophagy compared to DMSO control (Fig. 4E). However, inhibition of the MEKK3-MEK5-ERK5 pathway by genetic or pharmacological tools did not mitigate CCCP-induced mitophagy either in the absence or presence of ectopic Parkin overexpression (Fig. 4F, 4G, and Sup. Fig. 2F-I). The combined data indicate the MEKK3-MEK5-ERK5 pathway specifically promotes lysosome-mediated mitochondrial degradation under basal conditions and is not required for nonspecific bulk autophagy, general lysosome function, or damage-induced mitophagy nor for restraint of mitochondrial biogenesis.

### MEKK3-MEK5-ERK5 Pathway is Required for Differentiation and Mitophagy in Erythroid Progenitors

Previous studies suggest that ERK5-deficient cells develop altered nucleotide metabolism, impairing erythropoiesis in mice (Angulo-Ibanez et al., 2015). Erythropoiesis involves an ordered acquisition and loss of key differentiation markers (Koulnis et al., 2011). Because erythroid progenitors must eliminate their mitochondria in order to fully mature, we evaluated whether ERK5-regulated mitochondrial degradation participates in erythrocyte maturation (Moras, Lefevre, & Ostuni, 2017). To investigate the possible role of the MEKK3-MEK5-ERK5 pathway in erythrocyte maturation, Ter119-negative erythroid progenitors were isolated from murine fetal livers and induced to differentiate by addition of erythropoietin ex-vivo. We followed the expression of two specific differentiation markers: CD71, the transferrin receptor, whose expression decreases as erythroblasts mature, and Ter119, an antigen expressed on cell surfaces of more mature erythroblasts (Koulnis et al., 2011). BIX02188 or vehicle control (DMSO) were present throughout the ex-vivo differentiation process. Pharmacological inhibition of MEK5 impaired erythroid differentiation as evidenced by reduced percentages of cells reaching the most advanced Ter119^+^CD71^low^ stage of differentiation (Fig. 5A). Notably, MEK5 inhibition significantly increased basal mitochondrial content at all stages of differentiation (Fig. 5B). As differentiation progressed, both BIX02188-treated and DMSO-treated cells lost mitochondrial content over time; however, mitochondrial content was always significantly higher in BIX02188-treated cells relative to DMSO-treated cells at each stage (Fig. 5B). Similar results were observed when ERK5 was inhibited using XMD8-92 (Fig. 6A and 6B). Erythroid differentiation was similarly inhibited by genetic depletion of MEK5 using shRNA (Sup. Fig. 3). Together, these data indicate that MEK5-ERK5 signaling is required for proper erythroid differentiation and promotes clearance of mitochondria.

**Figure 5:**
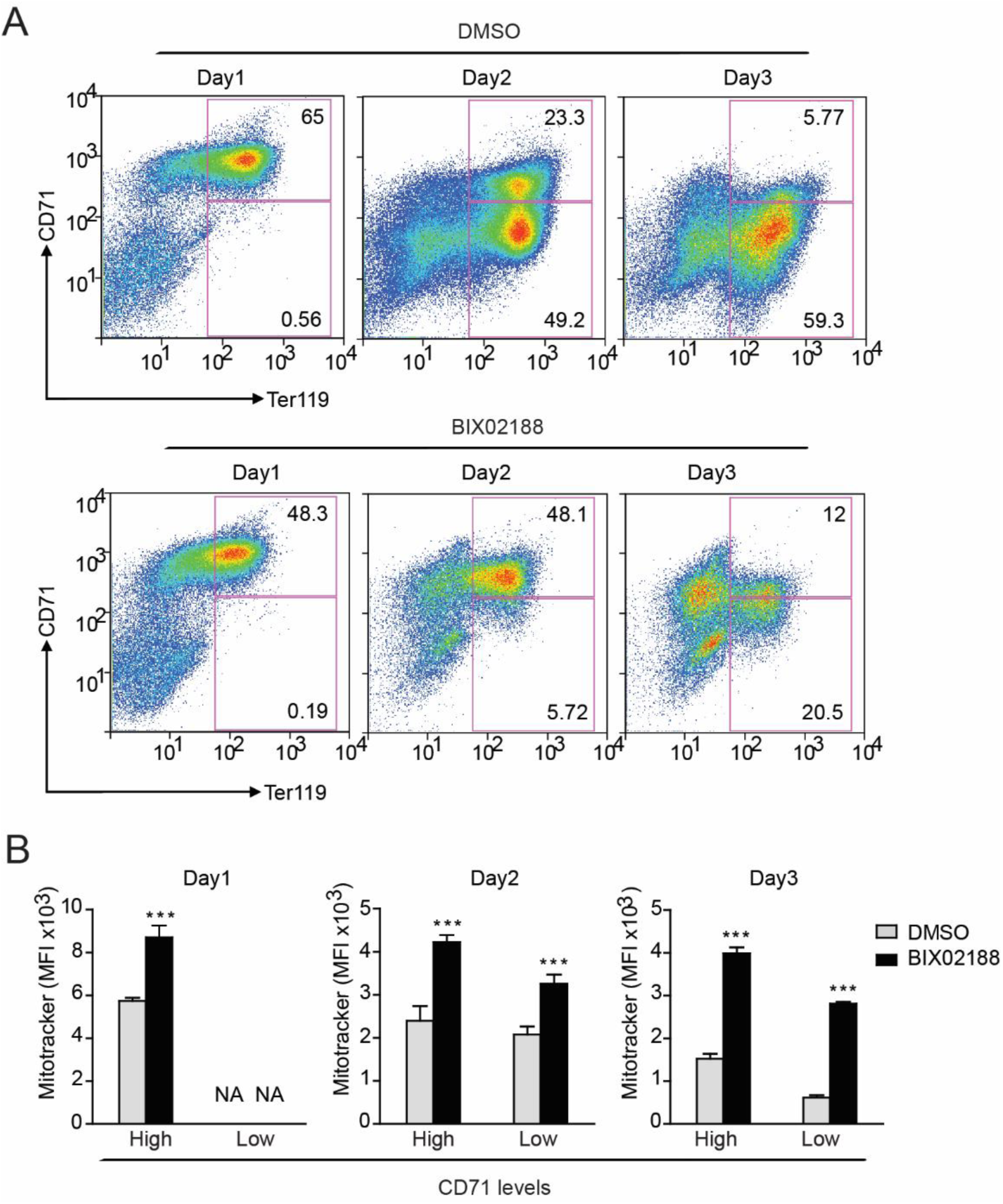
MEK5 is Required for Differentiation and Mitophagy in Erythroid Progenitors. (A) Pharmacological inhibition of MEK5 impairs erythroid differentiation. Murine erythroid progenitors were treated with the MEK5 inhibitor BIX02188 (30 μM) or vehicle control (DMSO) prior to initiation of differentiation. Representative profiles are shown. (B) Inhibition of MEK5 impairs mitochondrial clearance during erythroid differentiation. Mitochondrial mass was quantified in Ter119+CD71high and Ter119+CD71low erythroblasts using median fluorescence intensity (MFI) of MitoTracker Deep Red FM. The data represent three replicates. ***P < 0.001.

**Figure 6:**
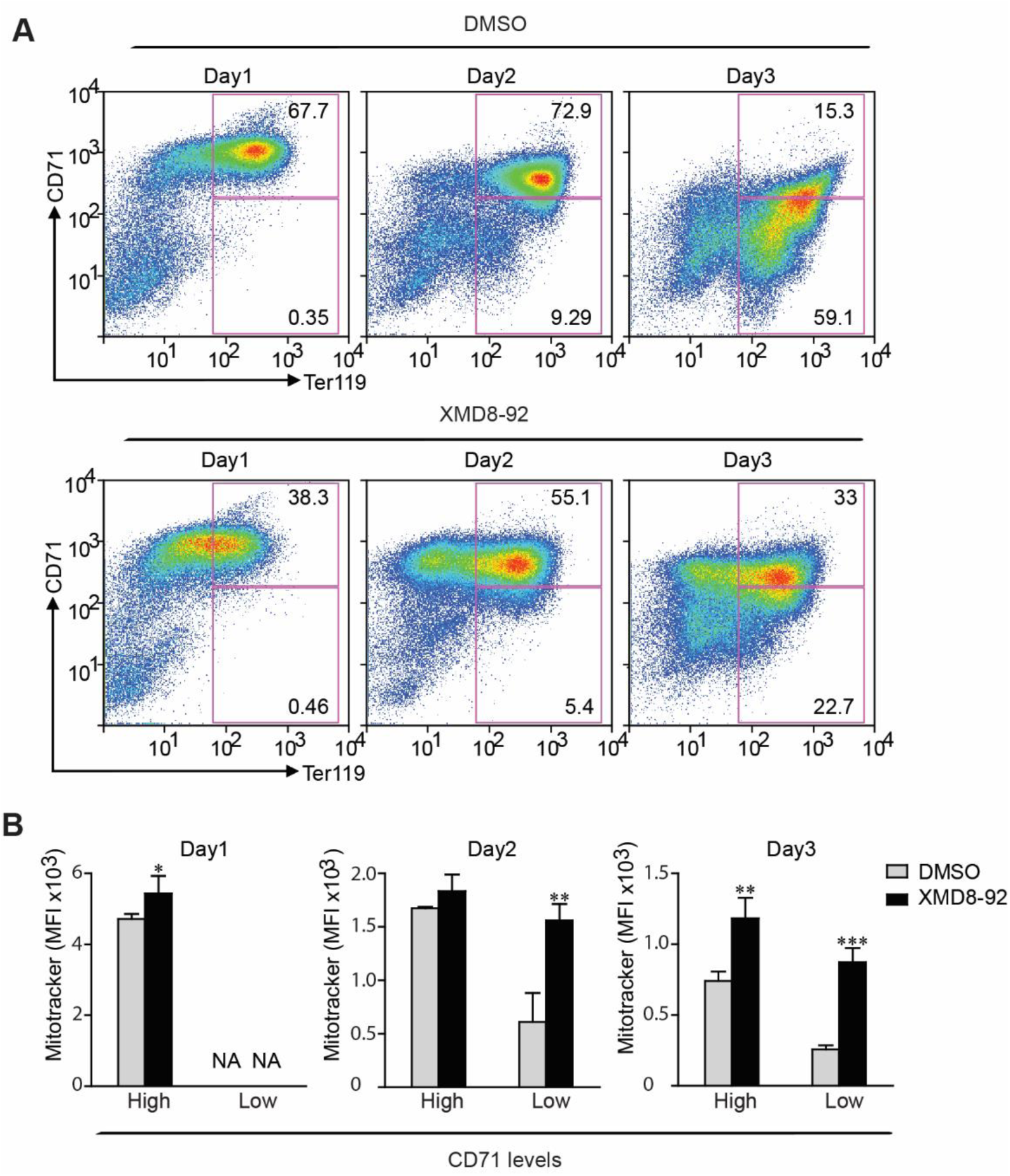
ERK5 is Required for Differentiation and Mitophagy in Erythroid Progenitors. (A) Murine erythroid progenitors were treated with the ERK5 inhibitor XMD8-92 (5μM) or vehicle control (DMSO) prior to initiation of differentiation. Representative profiles are shown. (B) Inhibition of ERK5 impairs mitochondrial clearance during erythroid differentiation. Mitochondrial mass was quantified in Ter119+CD71high and Ter119+CD71low erythroblasts using median fluorescence intensity (MFI) of MitoTracker Deep Red FM. The data represent three replicates. * P <0.05; **P < 0.01; ***P < 0.001.

## Discussion

Currently, there is substantial interest in the role autophagy plays in maintaining normal physiology and how dysfunctional mitophagy can lead to or enhance diseases. Our discovery of the MEKK3-MEK5-ERK5 pathway as a novel mitophagy regulator could facilitate a deeper understanding of how different mechanisms act to maintain homeostasis and prevent disease in different tissues. Using genetic and pharmacological inhibition of the MEKK3-MEK5-ERK5 pathway, we observed mitochondrial content increase in human cancer cells, mouse embryonic fibroblasts, and primary erythroid progenitor cells isolated from the murine fetal liver. This suggests that MEKK3-MEK5-ERK5 restrains mitochondrial accumulation in diverse cells and tissues. We also observed an increase in the levels of mitophagy receptor protein p62 after pharmacological or genetic inhibition of MEKK3-MEK5-ERK5 signaling. We confirmed that inhibition of pathway activity results in decreased lysosome-dependent degradation of mitochondria using mitochondrial targeted mKeima, without affecting lysosome acidification or non-selective bulk-autophagy levels in the cell. Taken together, we found that the MEKK3-MEK5-ERK5 pathway promotes mitophagy without the addition of exogenous mitochondrial damage, which could have great implication on studying mitophagy in the context of development and disease.

Additionally, our findings provide evidence for the importance of p62 in mitophagy under basal conditions. We have found that SQSTM1/p62 routinely delivers mitochondria to lysosomes without the addition of exogenous damage. We have found that inhibition of autophagy non-specifically with Bafilomycin A1 increases total mitochondrial load in cells, indicating that mitophagy is an ongoing maintenance process. Whether MEKK3-MEK5-ERK5 signaling controls mitochondrial degradation by directly or indirectly regulating p62 remains to be determined, although we have ruled out the possibility that MEKK3-MEK5-ERK5 signaling is required for p62 expression or stability. Direct interactions between the PB1 domain of p62 and the PB1 domains of MEKK3 and MEK5 have been reported, providing a potential means for direct regulation of p62 by the MEKK3-MEK5-ERK5 kinase cascade (Lamark et al., 2003; Nakamura et al., 2010).

We found that basal mitochondrial degradation, which requires p62 and MEKK3-MEK5-ERK5 signaling, is quite distinct from damage-induced mitophagy, which does not require p62 or MEKK3-MEK5-ERK5. We sought to investigate whether developmentally programmed mitochondrial degradation in the erythrocyte lineage required MEK5-ERK5 signaling as a well-established model of mitochondrial clearance in the absence of exogenous damage. Although impaired maturation of erythrocytes upon MEK5 or ERK5 inhibition precluded full assessment of the impact on mitochondrial turnover, the mitochondrial content was increased at each stage of erythroid maturation in MEK5-ERK5 inhibited cells. Taken together with the lack of a requirement for MEKK3-MEK5-ERK5 signaling in CCCP-induced mitophagy, it appears that the MEKK3-MEK5-ERK5 pathway may be solely designated for basal mitochondrial clearance. It is also possible that MEKK3-MEK5-ERK5 signaling may be required for induction of mitochondrial degradation by other stimuli, such as statin treatment, hypoxia, or forced metabolic switch from glycolytic metabolism to oxidative phosphorylation (Andres et al., 2014; L. Liu et al., 2012; Sin et al., 2016; Wilkinson, Sidaway, & Cross, 2018). These questions remain to be addressed in future work.

While MEKK3-MEK5-ERK5 signaling has been implicated in many aspects of mammalian development and disease, it is also the most recently discovered MAP kinase pathway, and much work needs to be done to fully understand how the MEKK3-MEK5-ERK5 kinase cascade is acting. It will be interesting to determine whether any of the previously identified roles for the ERK5 pathway in development, physiology, or disease represent an underlying role for mitochondrial degradation. For example, ERK5 signaling is well established to play an anti-apoptotic role, and mitochondria harbor several pro-apoptotic molecules including cytochrome *c*, Smac/DIABLO, and AIF (Adrain, Creagh, & Martin, 2001; Baechler, Bloemberg, & Quadrilatero, 2019; Nithianandarajah-Jones et al., 2012). It is possible that active ERK5 signaling antagonizes autophagy in part through elimination of pro-apoptotic molecules via delivery of mitochondria to lysosomes for degradation. These and other interesting hypotheses arising from our current work remain to be tested in future studies.

## Acknowledgements

We would like to thank M. A. White, F. Rivas and M. Kundu for valuable discussions. We are grateful to H. Gan and M. Wang for their assistance. We wish to acknowledge support from the Center for Advanced Genome Engineering, Flow Cytometry and Cell Sorting Facility, Cell and Tissue Imaging Center, and Hartwell Center for Biotechnology. This study was supported in part by the National Institute of General Medical Sciences of the National Institutes of Health under Award Number R01 GM132231 (M.B.P.), National Natural Science Foundation of China 81873430 (Z.H.), the National Cancer Institute of the National Institutes of Health under Award Number P30 CA021765 (Charles Roberts), the American Lebanese Syrian Associated Charities (ALSAC), and St. Jude Children’s Research Hospital.

## Conflict of Interest

The authors declare no conflicts of interest.

## Supplemental Information

**Supplemental Figure 1:**
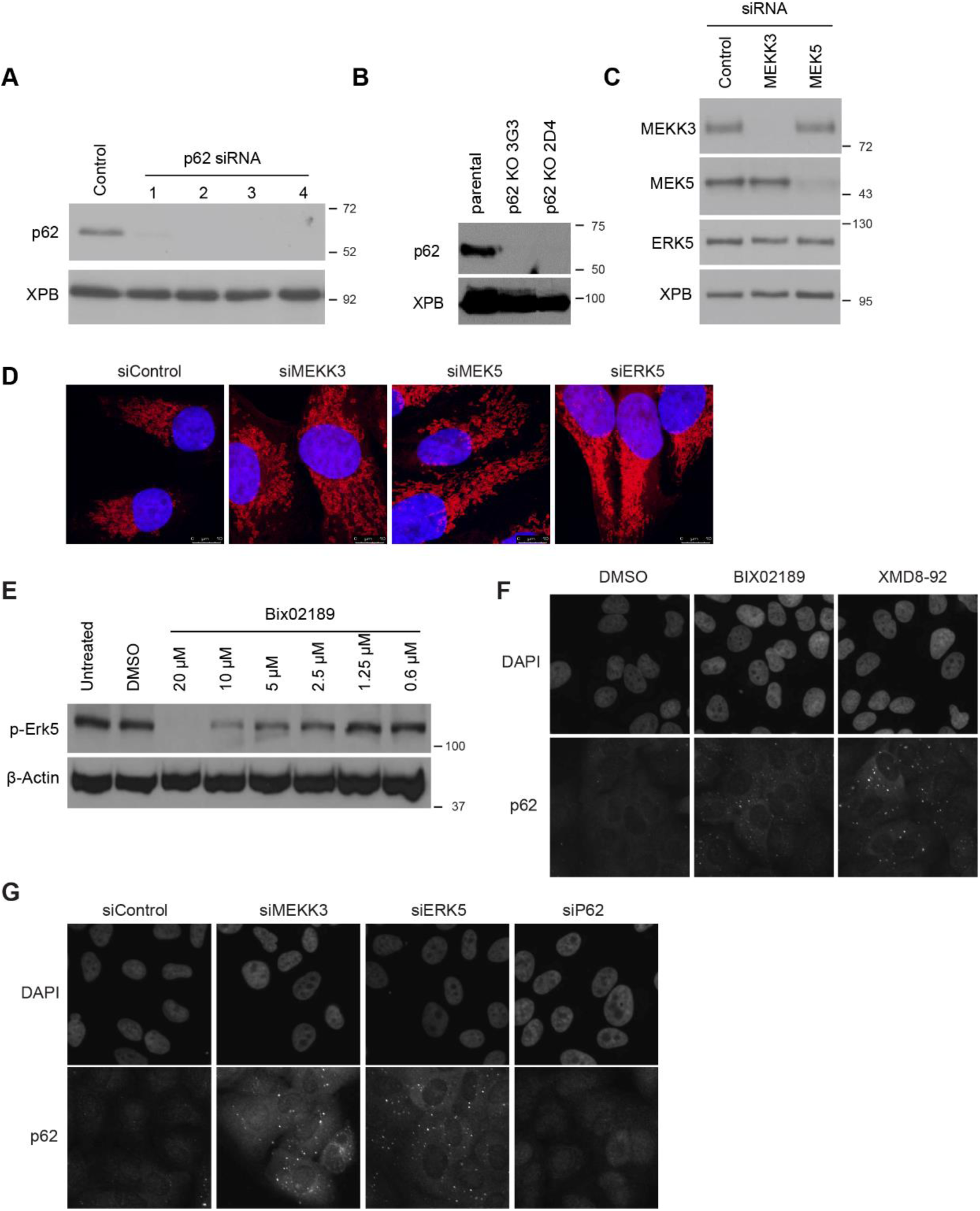
The MEKK3-MEK5-ERK5 pathway prevents accumulation of excess mitochondria. (A) U2OS cells were transfected with the indicated siRNAs. 72 hours later lysates were collected and analyzed by western blot using the indicated antibodies. (B) Parental and SQSTM1/p62 knockout U2OS cell lysates were collected and analyzed by western blot using the indicated antibodies. (C) U2OS cells were transfected with the indicated siRNAs. 72 hours later lysates were collected and analyzed by western blot using the indicated antibodies. (D) U2OS mito-mCherry cells were transfected with the indicated siRNA oligos. 72 hours later cells were fixed and imaged in z-stacks. Representative maximum intensity projections are shown. (E) Mouse embryonic fibroblasts were treated with vehicle (DMSO) or the indicated concentration of the MEK5 inhibitor BIX02189 overnight. (F, G) U2OS cells were either transfected with the indicated siRNAs for 72 hours or treated with the indicated drug at 10 µM overnight. Cells were fixed and stained with p62 antibody, and then imaged.

**Supplemental Figure 2:**
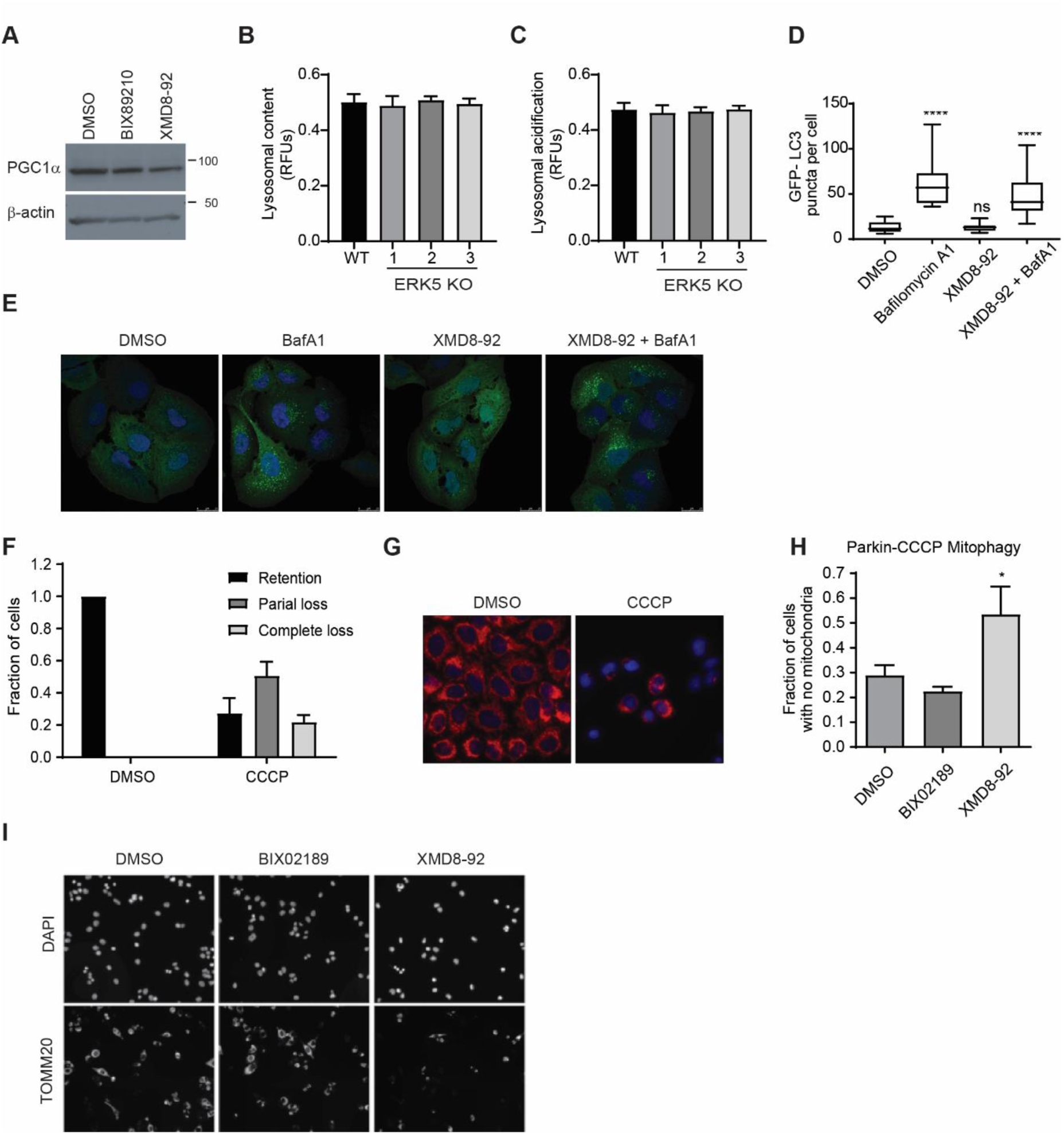
The MEKK3-MEK5-ERK5 pathway does not inhibit mitochondrial biogenesis or drive bulk autophagy. (A) U2OS cells were treated with vehicle (DMSO) or 10 μM of the indicated compound overnight. Lysates were collected and analyzed by western blot with the indicated antibodies. (B, C) WT and ERK5 KO U2OS cells were plated in a 96 well plate. Lysosomal content and acidification was measured by staining with LysoTracker Red and LysoSensor Green respectively. Cells were then imaged and relative fluorescence was quantified. (D, E) U2OS GFP-LC3B cells were treated with DMSO, XMD8-92 (10 μM), Bafilomycin A1 (50 nM), or XMD8-92 + Bafilomycin A1 for two hours, fixed, stained with DAPI (blue), and imaged to assess the subcellular localization of GFP-LC3B. Representative images are shown in (E) and quantitation of the number of GFP-LC3B-positive puncta per cell from 19-21 cells per condition is shown in (D). (F, G) HeLa YFP-Parkin stable cells were treated with either vehicle control (DMSO) or 10 µM CCCP overnight. Cells were fixed, stained with anti-Tom20 (red) and DAPI (blue), and imaged. Individual cells were categorized using Tom20 fluorescence signal as exhibiting either mitochondrial retention, partial loss, or complete loss. Quantitation is shown in F and representative images in G. (H,I) HeLa YFP-Parkin stable cells were co-treated with 10 µM CCCP and vehicle control, BIX02189, or XMD8-92 overnight. Cells were fixed, stained with anti-Tom20 and DAPI, and imaged. The fraction of individual cells exhibiting complete mitochondrial loss under each condition is quantified in H. Representative images are shown in I.

**Supplemental Figure 3:**
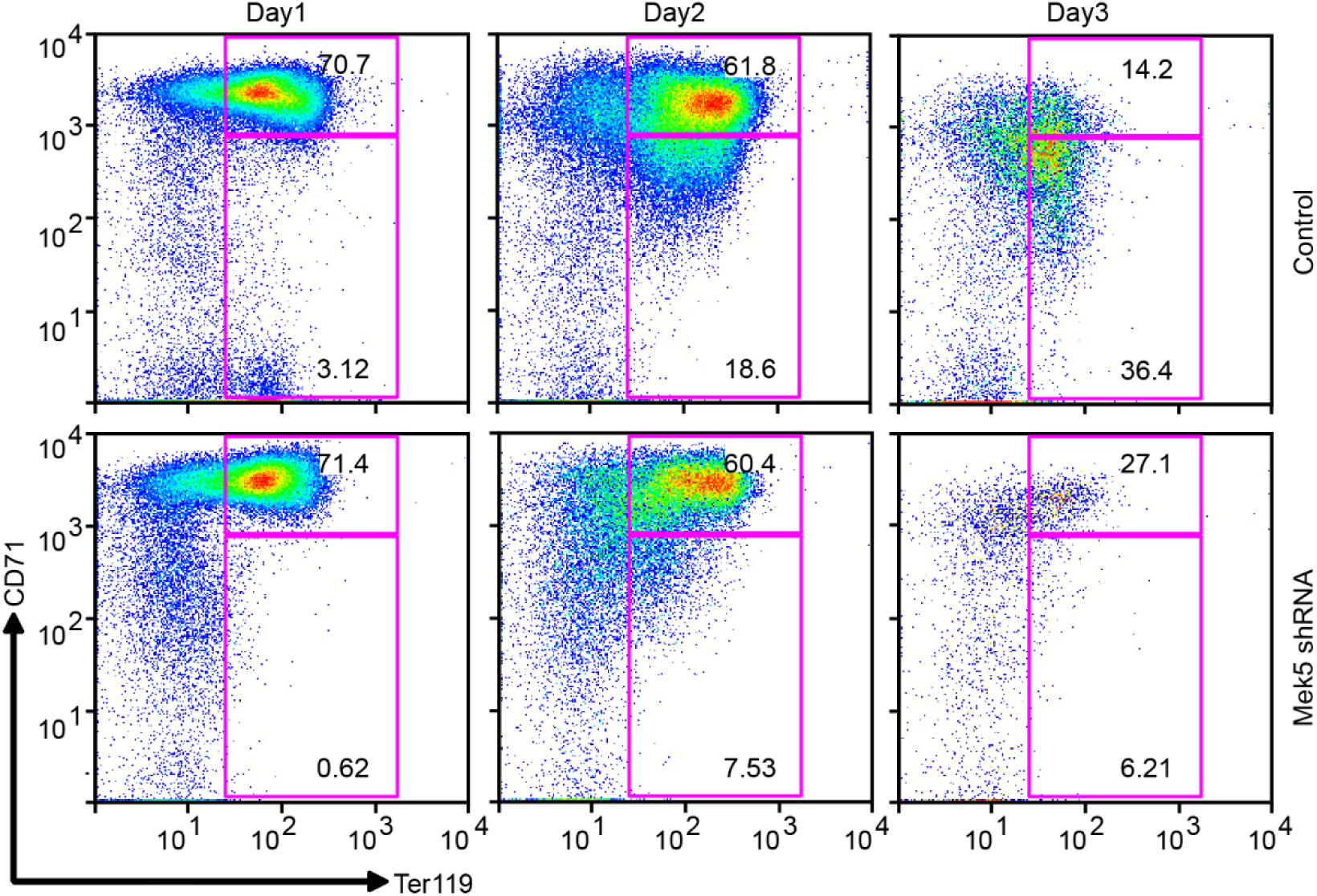
Inhibition of MEK5 impairs erythroid differentiation. MEK5 knockdown impairs differentiation of murine erythroid progenitors. Erythroid progenitors were transduced with vectors expressing shRNA for MEK5 or control shRNA prior to erythroid differentiation. Stages of differentiation were identified by CD71 and Ter119 staining via flow cytometry. Transduced cells, identified by GFP expression, were gated for analyses. Representative flow cytometric profiles are shown.

## Notes

### Competing Interest Statement

The authors have declared no competing interest.

